# “The ship is going down and we are powerless”: *The Impact of Federal Funding Changes on Researchers Training the Next Generation of Scientists*

**DOI:** 10.1101/2025.08.22.671828

**Authors:** Emily Mastej, Tiffany Do, Arghavan Salles

## Abstract

**Introduction:** Historically, federal grant terminations have been rare and almost exclusively used in cases of misconduct. Under the current presidential administration, however, grant terminations have been common, with thousands of grants terminated across federal agencies such as the National Institutes of Health (NIH) and the National Science Foundation. Although there have been scattered reports of the impact of these terminations on individual researchers, there has not yet been a systematic investigation of the impact on the overall scientific community.

**Methods:** As part of a survey for a larger study, we asked researchers involved in T32 training programs how likely they would be to go into science if they were a graduate student now (Likert-type item) and how federal funding changes have affected them and their labs (open-ended). We report mean and standard deviation for the Likert-type item (1=Not at all likely, 5=Extremely likely), and we performed inductive thematic analysis to analyze the open-ended responses. Two independent raters coded responses, developed a codebook, and applied the codebook to all responses. The interrater reliability was high, κ=0.84.

**Results:** Of the 490 researchers invited to participate in the survey, 277 (response rate 56.5%) completed it in full. Participants were relatively unlikely to choose to pursue science again, with only about one-third being somewhat or more likely to do so. Two hundred twenty-three people responded to the open-ended item about how federal funding changes have affected them. Coding resulted in 48 unique codes which fell into four themes: Impact on Finances, Impact on Trainees, Impact on Science, and Well-being. Participants described instability in their labs, concern for trainees’ well-being and careers, concern about the future of the entire scientific enterprise, and negative impacts on their and others’ well-being.

**Discussion:** Our data demonstrates the significant, overwhelmingly negative impact of the changes in federal funding on researchers involved in training the next generation of scientists. Not only are they concerned about how they can continue to do the work they have dedicated their lives to, but they are also worried about the future of science in the United States. This latter concern stems not only from the decreasing availability and reliability of federal funding but also due to the impact all this is having on trainees’ ability to do their research and find jobs where they can do science. These data are a call to action to our federal agencies, institutions, other agencies, and donors who understand the value of science to support researchers through these uncertain times. Otherwise, what has taken decades to build will be eliminated by a few anti-scientific decisions.

## INTRODUCTION

The arrival of the new presidential administration this year has ushered in significant and widespread changes to the operations of federal agencies that fund scientific research, such as the National Institutes of Health (NIH) and the National Science Foundation. These changes have led to widespread layoffs,^1^ which have impacted program officers and other key personnel^2^ responsible for administering grants and have caused significant delays in grant review processes. Additionally, thousands of grants have been outright terminated.^3^ Such terminations were previously rare and typically reserved for cases involving research misconduct or other serious violations.

Although a handful of articles have begun to address these funding changes,^4,5^ a substantial gap remains in our understanding of their long-term impact, particularly on the future of research and the career trajectories of early-stage investigators. As part of a larger research study on sexual harassment which, notably, was recently terminated by the NIH, we surveyed researchers involved in T32 training programs^6^ to understand how recent shifts in federal funding have impacted them, their work, and their labs. T32 programs play an important role in training the next generation of scientists by supporting both predoctoral and postdoctoral research training. Here we share the perspectives of T32-affiliated researchers and the impact of ongoing funding instability on them and their work.

## METHODS

Between May 8 and July 15, 2025, we conducted a cross-sectional survey of NIH T32 training program faculty who had previously enrolled in our sexual harassment study. We invited the faculty to fill out optional survey questions to further understand how recent federal funding changes have impacted the scientific community. The survey consisted of two key items. The first was a closed-end question asking participants if they were a graduate student today, how likely they would be, on a scale from 1 to 5 where 1 is ‘Not at all Likely’ and 5 is ‘Extremely Likely,’ to pursue a career in science. The second item was an open-ended question inviting respondents to describe how the recent funding changes have impacted them, their trainees, and their labs. The first item quantitatively assessed participants’ interest in a scientific career given the current funding changes and political climate. The second item qualitatively assessed participants’ personal experiences. The survey was distributed via email, and responses were collected using Qualtrics. All responses were confidential.

To assess the Likert-scale item, we calculated the mean and standard deviation of the responses. Analyses were conducted in R (R version 4.4.1). To examine the open-ended qualitative response, we used inductive thematic analysis, guided by a post-positivist framework. This approach acknowledges the existence of an external reality while recognizing the influence of researcher interpretation. Two independent coders (TD and AS) reviewed the data and developed an initial codebook through iterative discussion. Coding was conducted using Dedoose (Dedoose version 10.0.35), and discrepancies were resolved through discussion. After the initial codebook development, the coders coded all responses, with coding segments being each unique idea. Interrater reliability was assessed using Cohen’s Kappa and was high, 0.84. Final themes were identified collaboratively to capture the diversity of participant perspectives. The study team includes experienced researchers, a senior statistician, and a research assistant who have all been affected to some degree by the termination of the NIH grant we worked on collaboratively.

## RESULTS

### Study Population

Of the 490 faculty members invited to participate in the survey, 277 completed it (response rate 56.5%). Respondents’ demographic data are shown in Table 1. Gender, race and ethnicity, sexual orientation, and geographical location did not differ significantly between those who responded to the survey and those who elected to answer the optional open-ended question regarding federal funding.

**Table 1.** Demographic data of participants.^a,b^.

|  | Overall Cohort<br>N (%) | Respondents to<br>Open-ended<br>Funding Question<br>Cohort<sup>c</sup><br>N (%) |
| --- | --- | --- |
| **Total** | 277 | 223 |
| **Gender** |  |  |
| Man | 129 (46.5) | 99 (44) |
| Woman | 144 (52) | 123 (55) |
| Other | <5 | <5 |
| Prefer Not to Say | <5 | <5 |
| **Race/Ethnicity** |  |  |
| Asian/Asian American | 41 (15) | 33 (15) |
| Black/ African American | 12 (4.3) | 11 (5) |
| Hispanic/ Latinx | 9 (3.2) | 7 (3) |
| Middle Eastern/ North African | 8 (3) | <5 |
| White | 195 (70) | 160 (72) |
| More than one race | 7 (2.5) | 6 (3) |
| Prefer Not to Say | 5 (1.8) | <5 |
| **Sexual Identity** |  |  |
| Straight/Heterosexual | 242 | 194 |
| Gay or Lesbian | 9 | 7 |
| Bisexual | 8 | 7 |
| Asexual | <5 | <5 |
| Other | 6 | 6 |
| Prefer Not to Say | 7 | 5 |
| Unknown | <5 | <5 |
| **Region** |  |  |
| Midwest | 57 | 50 |
| Northeast | 65 | 49 |
| South | 102 | 82 |
| West | 53 | 42 |

<sup>a</sup>Cells with fewer than 5 people are shown as "<5" to protect participants' identity
<sup>b</sup>Options for race, ethnicity, gender, and sexual identity have been compressed into fewer categories than on the original survey to protect participants' identity
<sup>c</sup>There were no statistically significant differences in the distribution of demographic groups between these two cohorts, all $p \geq 0.62$ .

### Likert-Type Survey Question

The average response to the question about likelihood of pursuing science was 2.38 (standard deviation = 1.2). As shown in Figure 1, only about one-third indicated they would be somewhat or more likely to choose science if they were a graduate student now.

**Figure 1.**
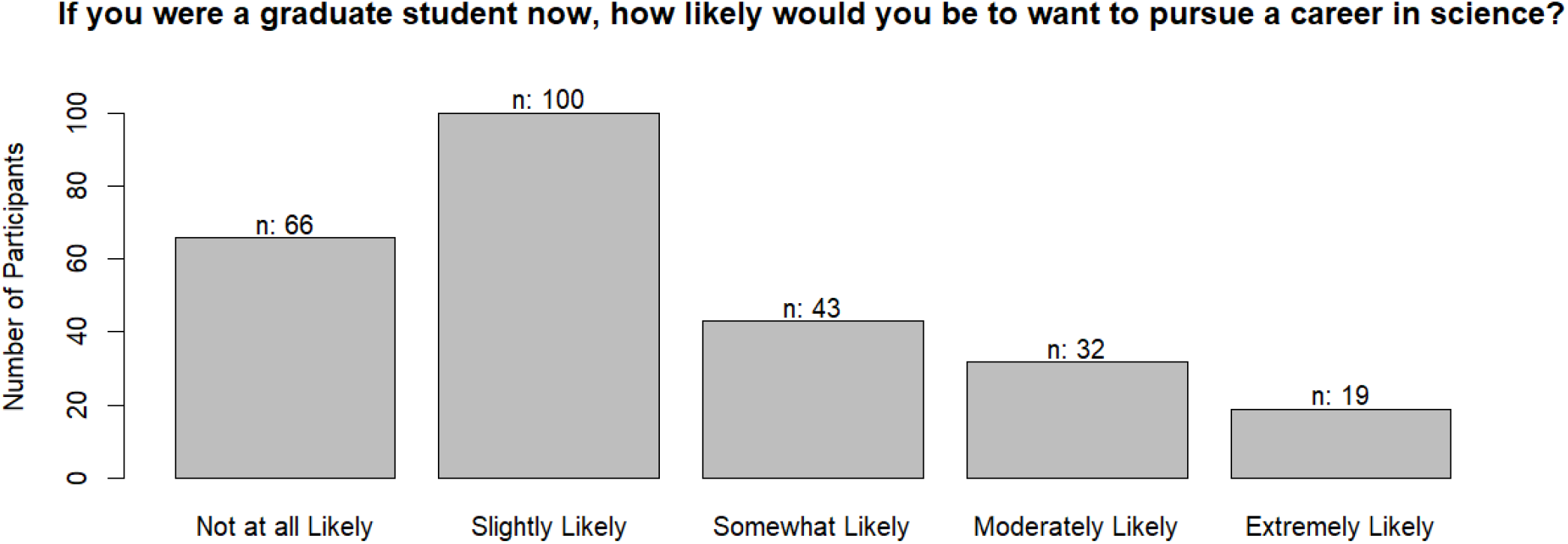
Likelihood of choosing a career in science again.

### Open-ended Survey Question

Two hundred twenty-three participants responded to an open-ended survey question that asked, “How have the changes in federal funding affected you and your lab?” When analyzing the open-ended responses, the coders identified 48 unique codes. Some codes were deemed non-substantive (eg, answers that simply said “yes”) and were excluded from further analysis. The substantive codes were further categorized into four themes to encompass participants’ responses: Impact on Finances, Impact on Trainees, Impact on Science, and Well-being. Illustrative examples of each theme can be found in Table 2.

**Table 2.**
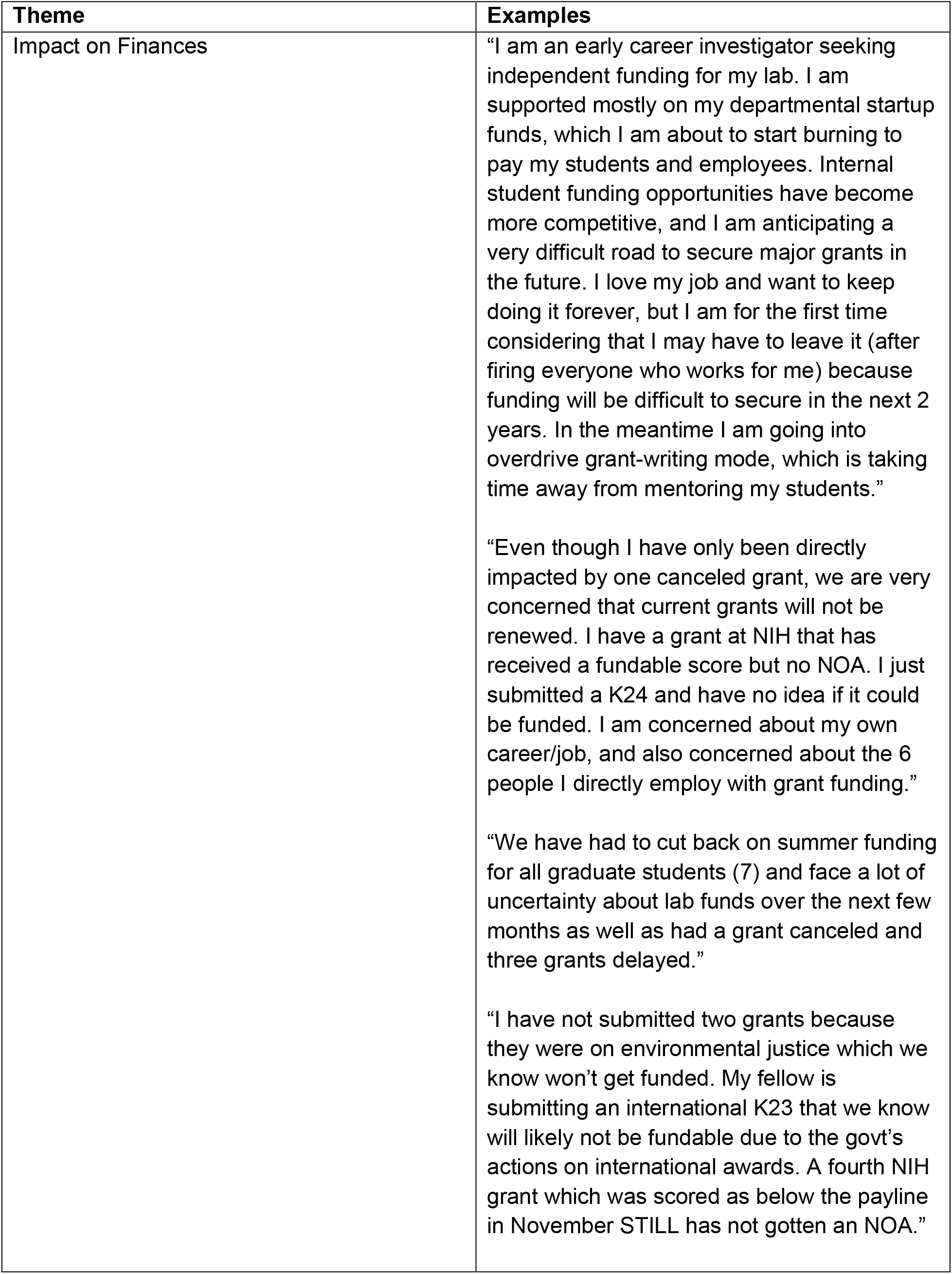
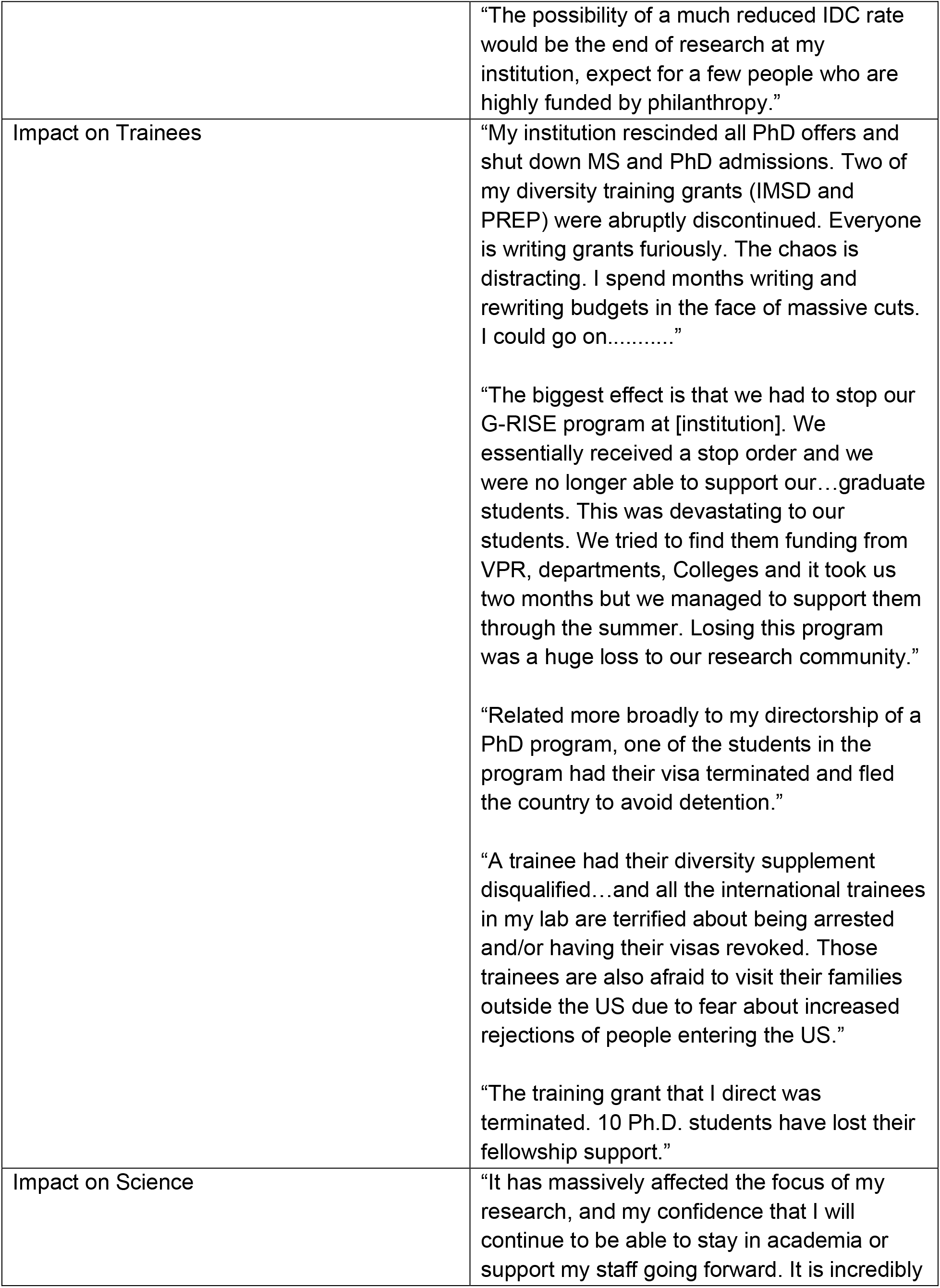
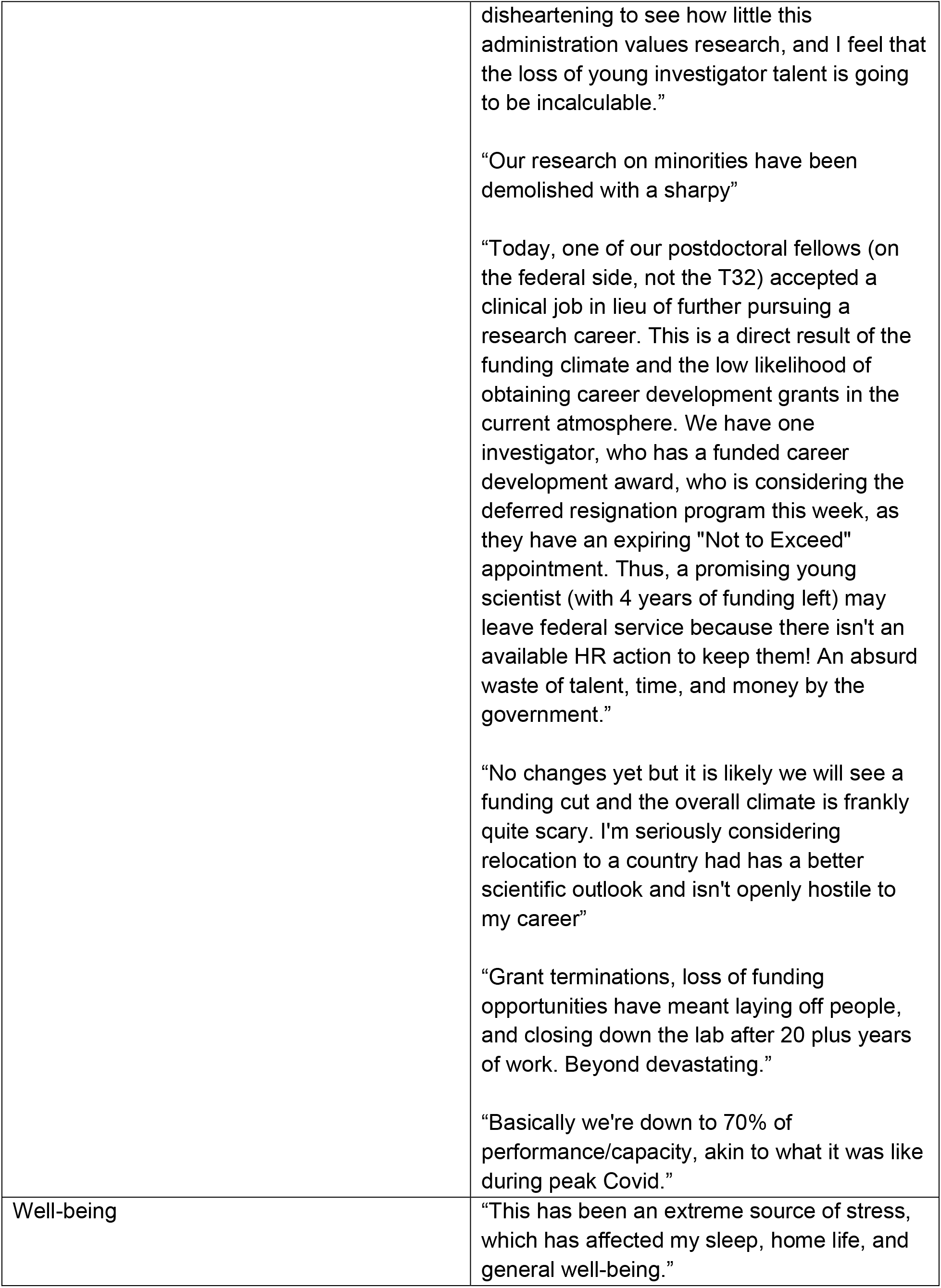
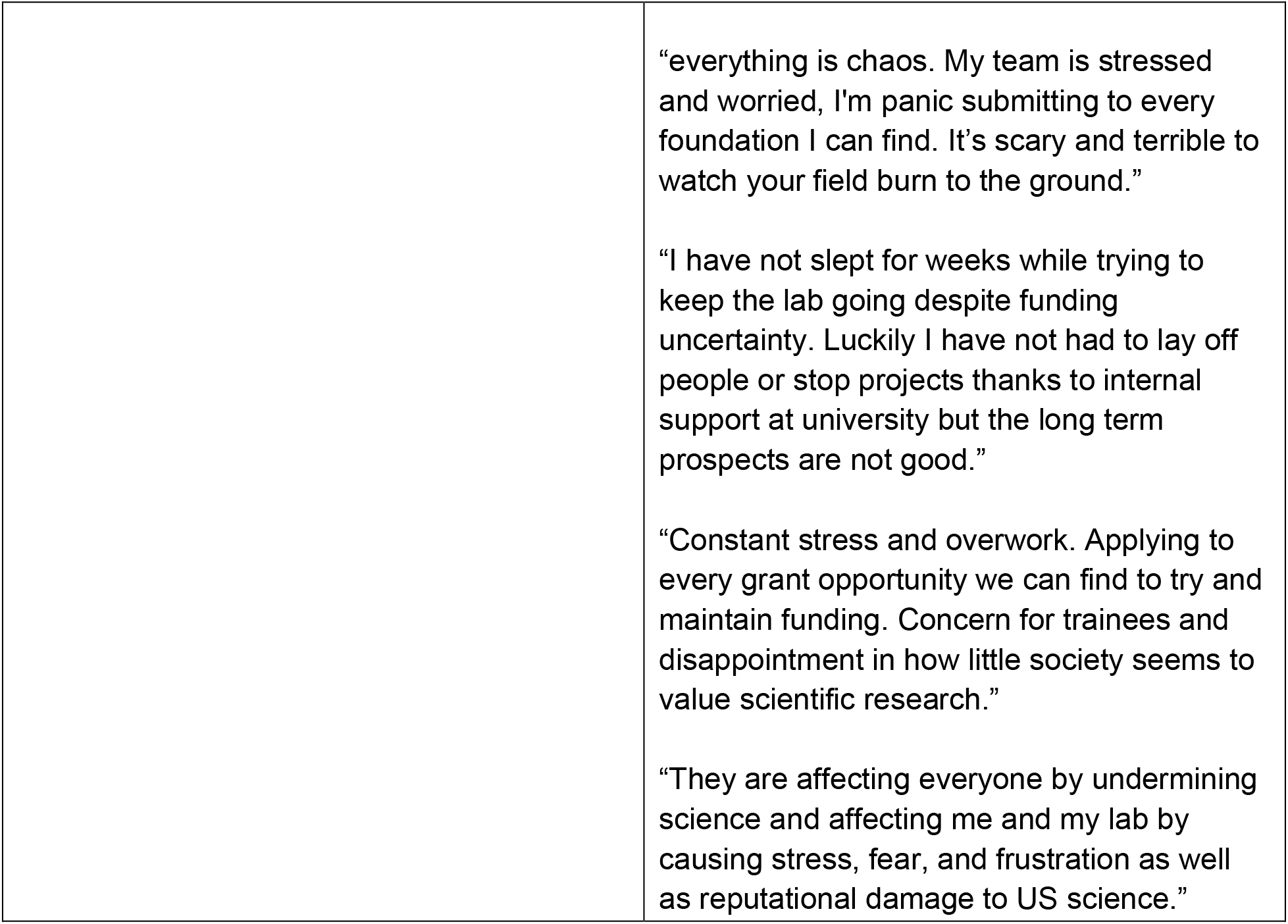
Qualitative themes and examples.

The theme of Impact on Finances encompasses participants’ references to their current financial insecurity due to funding changes. More specifically, participants described grant terminations, delays in funding agency reviews and funding notices, federal and local budget cuts, uncertainty about future funding, and potentially downsizing or even closing their labs. For example, one person wrote, “I am anticipating a very difficult road to secure major grants in the future. I love my job and want to keep doing it forever, but I am for the first time considering that I may have to leave it…because funding will be difficult to secure in the next 2 years.” Others echoed this concern: “I have money for now, but at some point, that will run out. The current funding agency I usually submit grants to is totally shutdown.

What will happen?”

The researchers invited to participate in this survey are all involved in training programs, and quite a few comments mentioned their decreased capacity to support trainees due to recent government policy changes. The theme of Impact on Trainees also includes concerns about training grant terminations, particularly those awarded to candidates from less represented backgrounds. Additionally, participants reported concerns about their trainees’ careers and well-being. One person wrote: “My students are having trouble finding jobs and are leaving research because of it. I am not recruiting new students as I don’t know if I can pay them…So soon I will have no students, because this is not how research works.” Many other comments mentioned reduction in trainee opportunities and graduate school admissions. One person wrote, “My institution rescinded all PhD offers and shut down MS and PhD admissions,” and another wrote, “The biggest effect is that we had to stop our G-RISE program at [institution]. We essentially received a stop order and we were no longer able to support our…graduate students. This was devastating to our students.” Others were concerned about the impact of immigration policies on trainees, with some even mentioning trainees having self-deported.

Many comments not only referred to Impact on Trainees but also related to Impact on Science, which encompasses broad concerns for attacks on science and the future of biomedical research. Participants wrote about having to change their research activities, often due to decrease in bandwidth to do research because of the need to do more grant writing or because their research topic no longer “effectuates agency priorities,” a phrase used in many of the termination notices^7^ sent to researchers. For example, one person wrote, “I am also concerned that this generation of graduate student will not have the opportunity to develop as postdocs and academics in the US which will lead to a drain of smart, diverse and creative people in science,” demonstrating their concern for their trainees and also the possible ‘brain-drain’ that could occur in the field of biomedical sciences. Many spoke of the negative impact on science independent of how these decisions affect trainees. For example, one person said, “The scientific enterprise is routinely being attacked and may come to a catastrophic end in the near future.” Yet another wrote, “We are all living in a climate of stress and uncertainty. The ship is going down and we are powerless.” Many of these comments included what we called a “sense of doom,” sentiments so ominous and gloomy we could not think of a better term to describe them. One comment read, “The past 5 months are by far the worst I’ve experienced in my science career (from grad student to professor) over the last 30+ years because we no longer know if there will be support for the work that we do, because science and research has been under attack and because this career that requires us to sacrifice so much in the pursuit of knowledge and truth has been so diminished in the eyes of the American people.”

The final theme that emerged from the data relates to Well-being, both that of the participants and those around them. Comments that were part of this theme related to anxiety, worry, changes to sleep, low morale, and other negative impacts on well-being. For example, one person wrote, “Morale is horrible. The uncertainty is driving anxiety to sky high levels.” Another wrote, “ongoing collective anxiety and anger…change the tenor of everyone’s daily work, that cost me sleep, and that make me more irritable, anxious, and unhappy.” Although not everyone in this study had been directly impacted by having a grant terminated, these comments showcase the feeling of overall anxiety, fear, hopelessness, and dread that currently fill many scientific researchers.

## DISCUSSION

Our data demonstrates the significant and overwhelmingly negative impact of the changes in federal funding on researchers involved in training the next generation of scientists. Relatively few researchers would choose a career in science again if they were in graduate school now. In their narrative comments, researchers described uncertainty in knowing how they would fund their lab, which caused extreme amounts of distress, even among those who had not yet lost any funding. They also were concerned about the future of their trainees and, more broadly and disturbingly, the future of scientific research in this country. Changes in immigration policies also were prominent in the responses, highlighting how changes unrelated to federal funding were impacting these researchers and their labs. Overall, the responses depict a time of great instability which is restricting scientists’ ability to do research, with multiple people either already having had to or considering shutting down their labs entirely. Given how recent all these changes have been, to our knowledge this is the first manuscript attempting to describe the scope of their impact.

There have been, however, scattered reports of the impact of grant terminations on specific individual researchers, and these largely align with our findings. For example, several researchers described to NPR the harm, including effects on their well-being, caused by their grants being terminated and the ensuing uncertainty in their careers.^8^ The New York Times profiled several early-career researchers whose grants had been terminated,^5^ demonstrating that the terminations impacted those working outside of the areas that apparently no longer aligned with agency priorities. Others have focused on the impact of grants to early-career researchers being terminated,^9^ demonstrating how the uncertainty makes it more challenging for young scientists to persist. Nature has also covered the additional challenges faced by international trainees.^10^ In many of the articles published on the topic, the attacks on science feature prominently, as in this piece published in the Harvard Crimson.^11^ On the whole, these reports align closely with our findings. However, our data extends beyond individual case reports to give an overview of what is happening across the country, not just to those whose grants have been terminated, but those who have previously earned federal funding for their work. This suggests the negative impacts of these destructive policies extend far beyond the thousands of individuals whose grants have been terminated to the entire scientific research community.

Indeed, the negative experiences shared with us were not restricted to those involved in studying topics that are no longer favored by the current administration, such as trans people,^12^ the broader LGBTQ+ community,^13^ climate change,^14^ COVID,^15^ HIV,^16^ misinformation and disinformation,^17^ workforce development,^18^ etc—and work that remains critical. Rather, among many researchers there was a sense of doom as they wondered whether they and their colleagues will be able to continue doing the work they’ve dedicated their lives to. The impact of these cuts also extends far beyond these researchers and their teams; data suggest NIH funding is critical to developing new treatments and cures.^19^ One review of medications approved by the revealed all but two of the 356 medications approved from 2010 to 2019 had been developed with funding from the NIH.^20^ In addition, NIH funding is a fantastic financial investment—every dollar spent yields over $2.50 in economic activity.^21^ Together with our findings, these data should serve as a call to action for federal agencies, institutions, other agencies, and donors who understand the value of science to support researchers through these uncertain times. Otherwise, what has taken decades to build will be eliminated by a few anti-scientific decisions.

